# Developmental control of E-cadherin junctions by mechanical contractility in *Drosophila* embryos

**DOI:** 10.1101/2024.08.29.610110

**Authors:** Na Zhang, Wangfang Liu, Huiying Lu, Matthias Häring, Fred Wolf, Jörg Großhans, Zhiyi Lv, Deqing Kong

## Abstract

Adherens junctions are very plastic structures that change their composition and structure depending on the state of the epithelial tissue, involving mechanical stress and tissue dynamics. In *Drosophila* embryos, adherens junctions mature only during gastrulation following their formation a few minutes before during cellularization. Although the developmental maturation is obvious during gastrulation, the factors and conditions controlling it have remained unclear. Here, we assay the levels, distribution pattern, and mobility of E-cadherin during *Drosophila* gastrulation. Indicating a maturation, we find an increase in total levels at junctions and a drop in the mobile fraction of E-cadherin molecules. Both developmental changes depend on Myo-II contractility. Consistent with stereotypic Myo-II regulation, we find that interference with Rho signaling leads to corresponding changes in E-cadherin levels and mobility, at least in part. Besides Rho signaling, Src42A contributes to junction maturation, as tested by both loss-function and gain-of-function situations. Together, we demonstrate developmental maturation of adherens junctions and identify Rho signaling and Src42A as upstream regulatory pathways.

## INTRODUCTION

Cell adhesion and its spatiotemporal regulatory process play critical roles in development and homeostasis ^1^. The classical cadherins are at the center of the adherens junction and, together with alpha-catenin, beta-catenin and further associated proteins, play an important role in tissue integrity, morphogenesis, and repair ^2^.

E-cadherin molecules generate cell-cell adhesion via Ca^2+^-dependent *trans*-interaction through their extracellular domains between adjacent cells. The intracellular C-terminal region forms a conserved complex with β-catenin and α-catenin that links E-cadherin to the cell cortex. This architecture and its adhesion function are evolutionarily conserved in all metazoans down to sponges ^3^. The mechanical forces generated by the contractile actomyosin cortex impinge on cell adhesion by altering the recruitments and molecular dynamics of E-cadherin at cell junctions ^4,5^.

Data from cultured mammalian cells and primary cells from zebrafish embryos support the idea that increased contractility results in increased recruitment or reduced turnover of E-cadherin at cell contact, what consequently expands the cell-cell contact ^6,7^. However, mechanical forces can affect E-cadherin also in an opposite manner. FRAP experiments in the MDCK cells demonstrated a correlation between increased tension and increased turnover of E-cadherin ^8^. VE-cad trans-interaction was stabilized by Rac1-mediated tension reduction in the endothelium ^9^. In addition, Src-induced de-regulation of E-cadherin depended on the phosphorylated-myosin activity and integrin signaling in colon cancer cells ^10,11^. These data point to a multifactorial and situation-dependent influence of cortical contractility on E-cadherin, yet the detailed mechanisms remain unexplored.

E-cadherin arranges in distinct submicron-scale clusters in the AJs ^3,12^. E-cadherin initial clustering is generated by cis-interactions, and the submicron-scale clustering is assumed to be dependent on the interaction between the E-cadherin-catenin complex and F-actin ^13–15^. Notably, FRAP experiments have demonstrated that E-cadherin turnover is slower for clustered than uniformly spread E-cadherin both in the cultured mammalian cells and *Drosophila* embryos ^6,16^. Furthermore, the immobilization of E-cadherin depends on F-actin and tension, probability due to a vital interaction between the E-cadherin-catenin complex and F-actin when under force ^17^. Clustering and immobilization is essential for E-cadherin-based mechanobiology ^4^. However, uncertainty remains about the function of clusters in AJs for the molecular dynamics of E-cadherin, especially in dynamic junctions.

During development, the AJs are dynamic and able to adapt to tissue morphogenesis while maintaining tissue integrity. Understanding the developmental control of E-cadherin serves as a model for the context-dependent plasticity of AJs. Here we focus on the ectodermal epidermis during germband extension in *Drosophila* embryos. *Drosophila* E-cadherin mutants (*shotgun*) ^18,19^ are characterized by various defects in distinct tissues during embryogenesis. This differential sensitivity may reflect the tissue activity, such that epithelia under mechanical challenge or dynamic epithelia, e. g. the lateral epidermis/germband during gastrulation, would require more E-cadherin protein to maintain tissue integrity ^20^. Junctions form during cellularization with the spot-like distribution of E-cadherin and are converted to proper belt-like AJs during gastrulation ^21,22^; meanwhile, more E-cadherin recruit associated molecules to the AJs ^13^. Further, it remains unclear if and how the location of E-cadherin clusters and molecular dynamics are regulated in the epidermis during gastrulation. Myo-II is activated by the Rho signaling and Src in the lateral epidermis, generating anisotropic contractility and driving the extension process of germband ^23–25^. It has remained unresolved, whether and how these components and mechanical challenges are involved in the developmental control of E-cadherin junctions during germband extension.

Here, we investigated the maturation of newly assembled adherens junctions shortly after cellularization to junctions in a mechanically challenged tissue undergoing remodeling. The gastrulating embryo provides insight into the dynamics of adherens junctions in relation to contractility given its stereotypic changes within a short period of time of about one hour. Using pharmacological and genetic interference we provide evidence for the control of developmental maturation of E-cadherin. We found that the decreased mobility of E-cadherin, distribution pattern and levels depend on the increased Myo-II dependent contractility. Consistent with genetic control of Myo-II, we found that Rho signaling and its upstream regulator RhoGEF2 as well as Src. Interestingly, both up and down regulation of Rho signaling and Src affect E-cadherin.

## RESULTS

### E-cadherin raising and immobilization during germband extension

Given the switch from syncytial to cellular development during cellularization (stage 5), the emergence of the first epithelium represents a starting point for when adherens junctions emerge. Soon after the epidermis is mechanically challenged during gastrulation (starting stage 6) and especially the lateral epidermis during germband extension (stage 7–9). As reported previously, E-cadherin initially forms spot AJs at the cellularization furrow during stage 5. Towards the end of cellularization, E-cadherin accumulates in a subapical region to form belt-like AJs characteristic of the epidermis ^21,22,26,27^ (Figures 1A–C).

**Figure 1.**
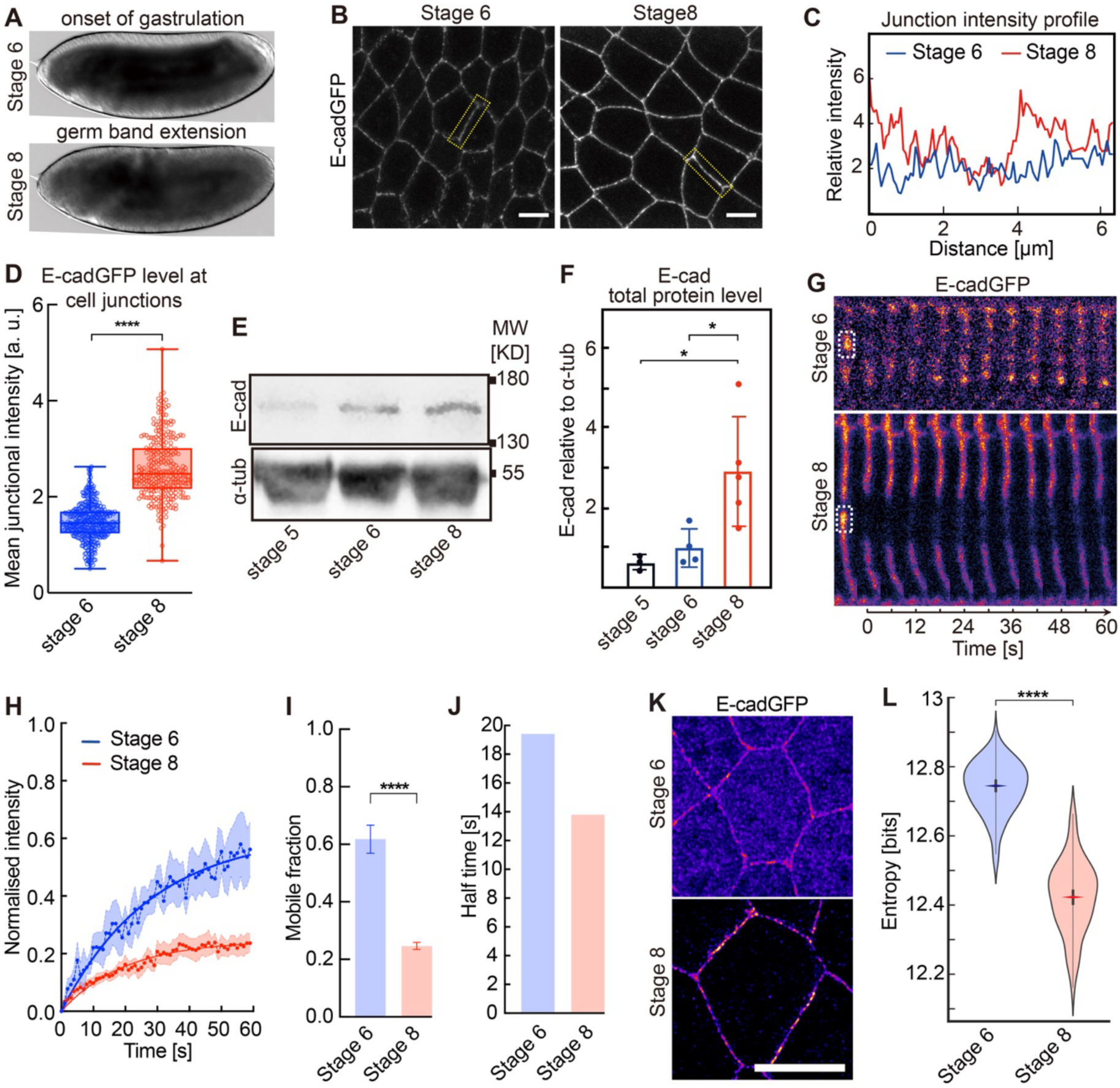
The developmental control of E-cadherin junctional raising and immobilization. **(A)** DIC images present the morphology change of *Drosophila* embryos from the onset of gastrulation (stage 6) to germ-band extension (stage 8). **(B)** Representative images of E-cadherin-GFP from the indicated stage embryos. **(C)** Intensity profiles of individual junctions from the boxes in (B). **(D)** Quantifications show increased E-cadherin recruitment at cell junctions from stage 6 to stage 8. N = 386 (stage 6) and 278 (stage 8) junctions from 5 embryos in each stage. **(E)** The western blot of total lysates with E-cadherin antibody has stages as indicated. Loading was controlled by western blot with α-tubulin antibody. **(F)** Quantification of E-cadherin protein level relative to α-tubulin from the western blot of total embryonic extracts, stages as indicated. The bar graph indicates the mean, the error bars indicate the SD, and each dot indicates one western blot. p-values were estimated by unpaired two-sided *t-test*, * p < 0.05. **(G–J)** E-cadherin-GFP fluorescence recovery after photobleaching in stage 6 versus stage 8 embryos. **(G)** Kymograph of E-cadherin-GFP pre- and post-bleaching, **(H)** normalized fluorescence intensity, **(I)** E-cadherin-GFP mobile fractions, and **(J)** mean recovery times after photobleaching. N = 6 (stage 6) and 10 (stage 8) junctions from more than three embryos in each stage. **(K)** Representative high-resolution images of E-cadherin-GFP by Airyscan jDeconvolution from stage 6 and 8 embryos. **(L)** Distribution of entropy values. The rhombus designates the median entropy value; thick bars indicate 95% confidence intervals, and the width is a kernel density estimate of the underlying distribution. Each entropy value was calculated by a 2.2 µm (200 pixels) long junction patch segment with a 0.44 µm (40 pixels) width. The boxes extend (in D and L) from the 25^th^ to the 75^th^ percentiles and the whiskers to the minimum and the maximum value with individual points. Data are mean ± SEM in (F and I), and solid curves indicate fitting in (F). Unpaired two-sided *t-test* estimates the p-values, **** p < 0.0001. Scale bars, 50 μm in (A), 5 µm in (B and K).

We first examined the levels and distribution of E-cadherin at junctions using a GFP tag introduced at the C-terminus of E-cadherin at the endogenous gene locus ^28^. By live imaging, we quantified E-cadherin-GFP levels at cell junctions (Figure 1B), finding an almost doubling of the mean intensities from stage 6 (onset of gastrulation) to stage 8 (germ-band extension) embryos, which is consistent with a previous report ^13^. We also examined total E-cadherin protein amounts by quantitative western blotting with extracts from manually staged embryos (Figure 1E). Consistent with the increased fluorescence at junctions, we measured a threefold increase in total E-cadherin protein levels from stage 5 to stage 8 (Figure 1F). Our data suggest that new E-cadherin molecules are assembled in adherens junctions during gastrulation.

E-cadherin forms stable clusters within adherens junctions ^3,12^, which we quantified by fluorescent recovery after photobleaching (FRAP) of E-cadherin-GFP. At the onset of gastrulation in stage 6, about 60% of E-cadherin molecules are exchanged within a minute. About 40 minutes later, in stage 8 embryos, the mobile fraction was reduced to about 20% (Figure 1G, H). By fitting to exponential functions, we calculated a significant decrease in the mobile fraction from 0.62±0.05 in stage 6 to 0.25±0.01 in stage 8 embryos (Figure 1I). The kinetic constant changed from 19 s half-time in stage 6 and 14 s in stage 8 embryos (Figure 1J).

Next, we developed an assay for the pattern of E-cadherin distribution along junctions. Fixed and stained embryos were imaged with AiryScan optics and subjected to joint deconvolution to achieve resolutions towards 100 nm. The E-cadherin clusters resolved in this way allowed us to calculate the entropy as a measure for their distribution. Entropy generalizes the notion of variance of distribution to arbitrary multi-dimensional and multi-modal distribution independent of intensity ^29,30^. Thereby, larger entropy values indicate a higher variability in the E-cadherin distribution and higher uncertainty about the location of E-cadherin clusters. We segmented the junctions and calculated the entropy values for junction segments of 2.2 µm x 0.44 µm (200 x 40 pixels). We detected a significant decrease in the entropy values from 12.740±0.01 bits (mean±95% confidence intervals) in stage 6 to 12.425±0.02 bits in stage 8 embryos (Figures 1K, L).

In summary, we established an assay for the developmental changes of adherens junctions as indicated by increasing levels and decreasing mobility and distributional variability of E-cadherin.

### Actomyosin contractility promotes junctions maturation

Actomyosin contractility promotes AJs stability and E-cadherin clustering in cultured cells ^31^, E-cadherin clusters’ stabilization in mammalian and zebrafish cells in culture ^6,7^, and in *Drosophila* embryos ^32^. We examined the non-muscle Myosin-II (Myo-II) by immunostaining using an antibody against Zipper, the myosin-heavy chain in *Drosophila* ^33,34^. During gastrulation, non-muscle Myosin-II obviously increased in the cytoplasm and at junctions from stage 6 to stage 8 (Figure S1A), consistent with previous reports ^35–37^. We quantified Myo-II by measuring the staining at the junctions excluding vertices (tricellular junctions), finding a significant increase from stage 6 to stage 8 embryos (Figure S1B).

As E-cadherin levels and mobility change in parallel to Myo-II levels, we tested their dependence by inhibiting actomyosin contractility with a chemical inhibitor (20 mM Y-27632) of the upstream kinase ROCK ^38^. Consistent with previous studies ^39^, we detected a loss of E-cadherin-GFP from cell junctions following injection of the ROCK inhibitor (Figure 2A). The mean intensity of E-cadherin-GFP decreased to less than half of the water-injected embryos (Figure 2B). For the E-cadherin molecules remaining at the junctions, we detected a large mobile fraction similar to the wild-type stage 6 embryos, as revealed by the FRAP experiments following the induction of the ROCK inhibitor. To ensure experimental consistency, the ROIs of bleaching were set with the same size in the wild-type embryos, even though E-cadherin clusters similar to those in wild-type embryos were rarely detected in the Y-27632-injected embryos. Representative kymographs of cell junctions and fluorescence curves show a clear effect of the ROCK inhibitor (Figures 2C and D). The mobile fraction of E-cadherin-GFP was a threefold increase in Y-27632-injected embryos compared to wild-type stage 8 embryos (Figure 2E). The half-time is 8 s in Y-27632-injected embryos, while it is 14 seconds in wild-type embryos (Figure 2F), suggesting a faster recovery due to Y-27632-injection. In summary, the effects of Myo-II inhibition indicate the increased actomyosin contractility from stage 6 to stage 8 contributes to the developmental control of adherens junction during gastrulation.

**Figure 2.**
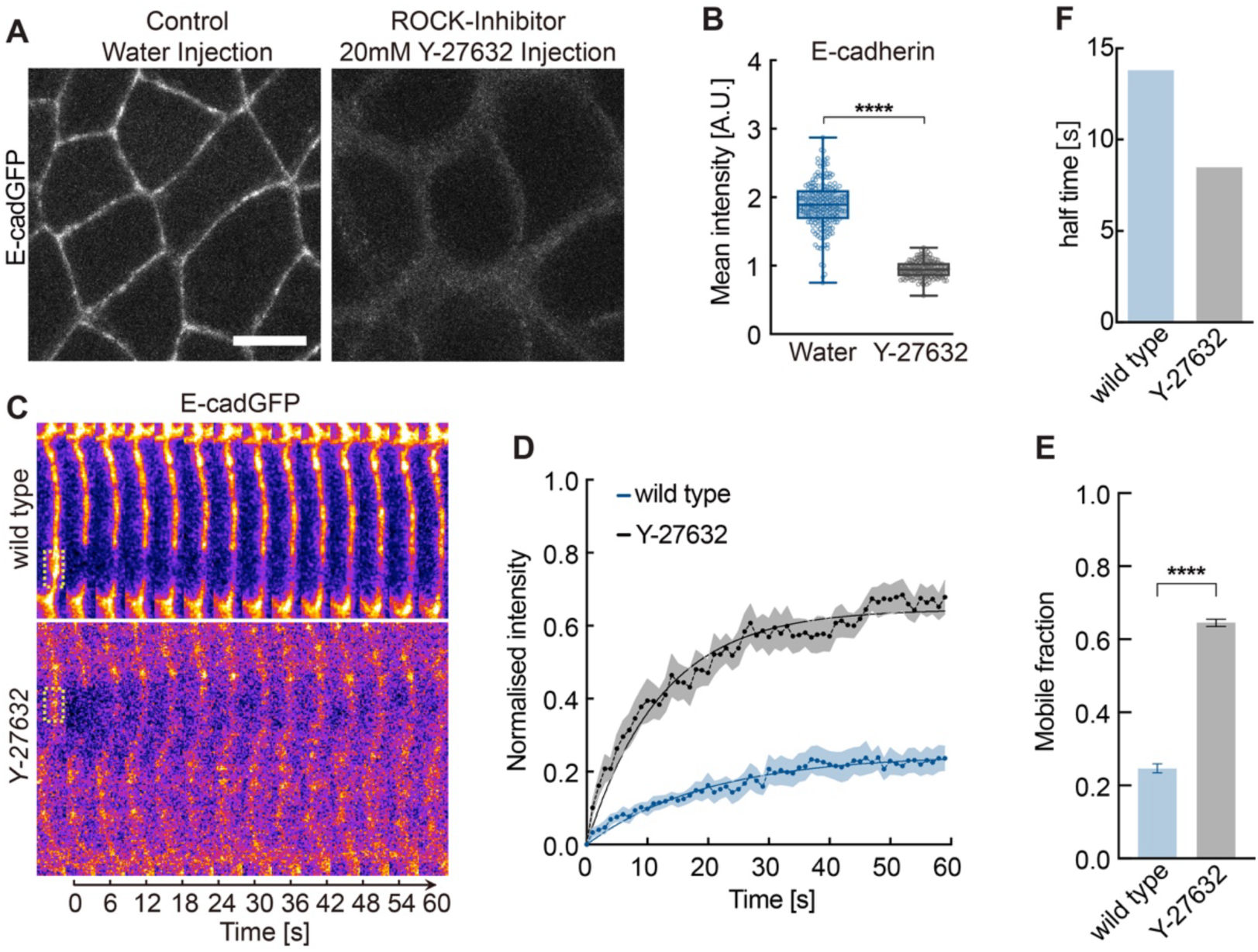
ROCK-inhibitor leads to junctional reduction and mobilization of E-cadherin. **(A)** Representative images of E-cadherin-GFP from water or ROCK inhibitor (20 mM Y-20mM Y-27632) injected embryos. **(B)** Quantifications show reduced junction E-cadherin recruitment from the ROCK inhibitor injected embryos at cell junction. N = 196 (water) and 146 (Y-27632) junctions in 4 water and ROCK inhibitor-injected embryos. **(C–F)** E-cad-GFP fluorescence recovery after photobleaching in wild type versus ROCK inhibitor injected embryos. **(C)** Kymograph of E-cadherin-GFP pre- and post-bleaching, **(D)** normalized fluorescence intensity, **(E)** E-cadherin-GFP mobile fractions, and **(F)** mean recovery times after photobleaching. N = 10 (wild type) and 11 (Y-27632) junctions in more than five wild-type and ROCK inhibitor-injected embryos. The boxes extend from the 25^th^ to 75^th^ percentiles, and the whiskers are at the minimum and maximum value with individual points in (B). Data are mean ± SEM in (D and E), and solid curves indicate fitting in (D). Unpaired two-sided *t-test* estimates the p-values, **** p < 0.0001. Scale bars 5 µm.

### Rho1 regulates E-cadherin junctional recruitment and immobilization

Previous studies have shown that Rho signaling actomyosin contractility through the activation of ROCK in the epidermis during gastrulation ^32,40,41^. To demonstrate a link between Rho signaling and developmental control of E-cadherin junctions, we expressed either a constitutively active version of Rho1 (Rho1 CA) or a dominant negative version of Rho1 (Rho1 DN) using the GAL4-UAS system ^42^. As expected, Myo-II fluorescence was increased by Rho1 CA expression while decreased by Rho1 DN in stage 8 embryos (Figure 3A). We employed a myosin regulatory light chain-mCherry transgene for quantification (Figures 3C and D, S2). Both junctional and medial Myo-II levels were reduced significantly in Rho1 DN embryos (Figures 3A, C, D), while increased junctional Myo-II levels were detected in Rho1 CA embryos, and medial Myo-II was not significantly affected (Figures 3B-D).

**Figure 3.**
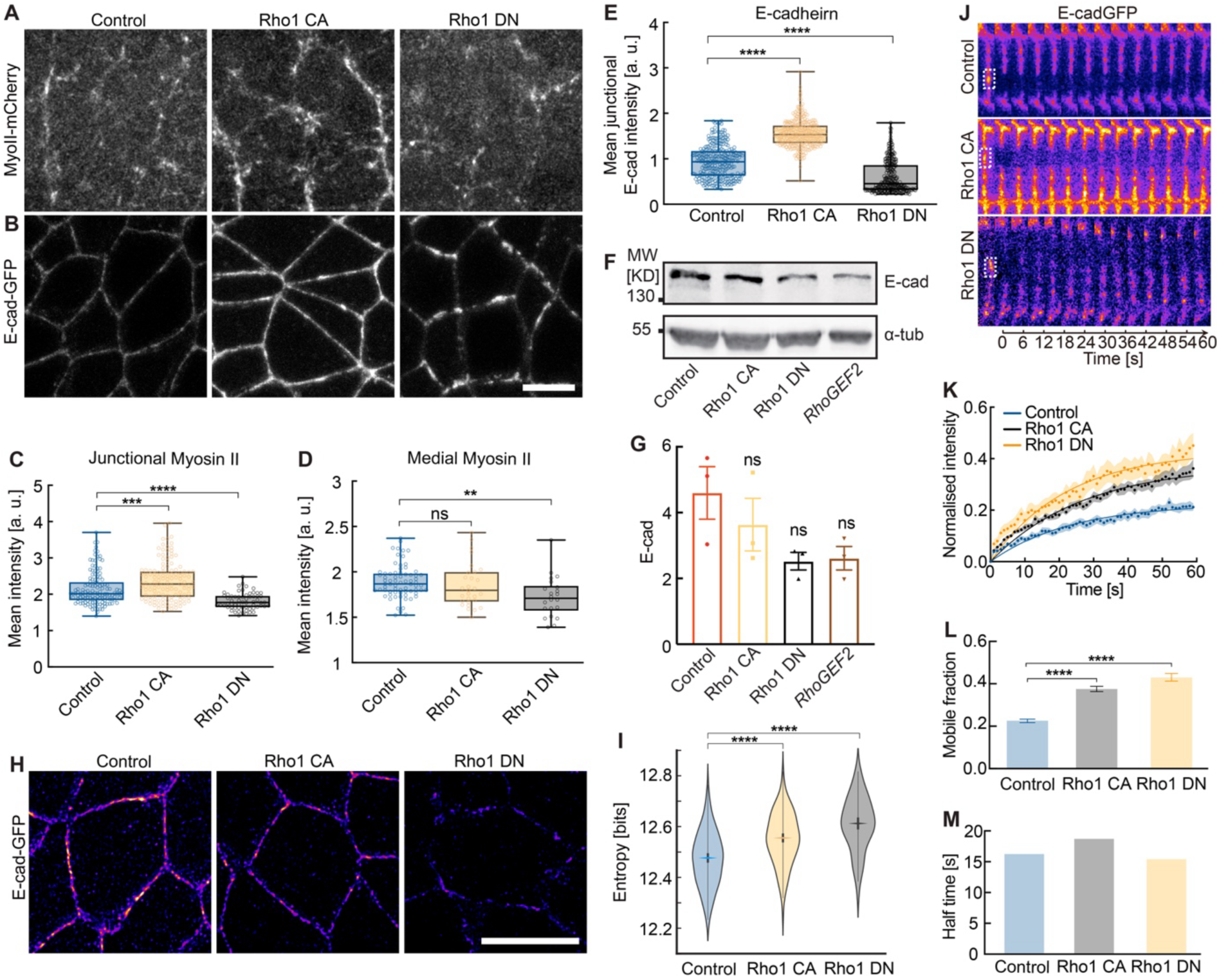
Rho1 regulates E-cadherin recruitment, distribution, and immobilization at cell junctions. **(A, B)** Representative images of Myosin-II-mCherry **(A)** and E-cadherin-GFP **(B)** from control, Rho1 CA (constitutively active Rho1), and Rho1 DN (dominant negative Rho1) embryos. **(C, D)** Quantifications show the junctional and medial Myosin-II were altered in Rho1 CA and Rho1 DN embryos. N = 133 (control), 131 (Rho1 CA), and 62 (Rho1 DN) junctions **(C)**; N = 71 (control), 29 (Rho1 CA), and 22 (Rho1 DN) cells **(D)** from more than 3 embryos in each genotype. **(E)** Quantifications show junction E-cadherin recruitment alters in Rho1 CA and Rho1 DN embryos. N = 298 (control), 321 (Rho1 CA), and 289 (Rho1 DN) junctions from 5 embryos in each genotype. **(F, G)** No significant difference in the E-cadherin total protein was detected from Rho1 mutants compared to control embryos. **(F)** Western blot of total embryonic extracts (3-6 h) with E-cadherin antibody, phenotypes as indicated. Loading was controlled by western blot with α-tubulin antibody. **(G)** Quantification of E-cadherin protein level relative to α-tubulin from the western blot of total embryonic extracts, phenotypes as indicated. The bar graph indicates the mean, the error bars indicate the SD and each dot indicates one western blot. P-values were estimated by unpaired two-sided *t-test*, ns > 0.05. **(H)** Representative high-resolution images of E-cadherin-GFP by Airyscan jDeconvolution from control, Rho1 CA, and Rho1 DN embryos. **(I)** Distribution of entropy values. The rhombus designates the median entropy value; thick bars indicate 95% confidence intervals, and the width is a kernel density estimate of the underlying distribution. Each entropy value was calculated by a 2.2 µm (200 pixels) long junction patch segment with a 0.44 µm (40 pixels) width. Scale bars, 5 μm. **(J-M)** E-cadherin-GFP fluorescence recovery after photobleaching in control versus Rho1 CA and Rho1 DN embryos. **(J)** Kymograph of E-cadherin-GFP pre- and post-bleaching, **(K)** normalized fluorescence intensity, **(L)** E-cadherin-GFP mobile fractions, and **(M)** mean recovery times after photobleaching. N = 16 (control), 15 (Rho1 CA), and 15 (Rho1 DN) junctions from more than five embryos in each genotype. The boxes extend from the 25^th^ to the 75^th^ percentiles and the whiskers to the minimum and the maximum value with individual dots in (C-E). Data are mean ± SEM in (K and L), and solid curves indicate fitting in (B). Unpaired two-sided *t-test* estimates the p-values, ns > 0.05, ** p < 0.01, *** p < 0.001, **** p < 0.0001.

Having established an experimental interference of Rho signaling and Myo-II levels, we assayed the consequences on E-cadherin levels, mobility and distribution pattern. Following increased or decreased Rho signaling, E-cadherin-GFP fluorescence at junctions changed correspondingly (Figures 3A-E), while total E-cadherin protein levels were not obviously affected, as assayed by western blot (Figures 3F and G). Rho signaling also controls the distribution of the E-cadherin clusters. Unexpectedly, larger entropy values were detected both in Rho1 CA and Rho1 DN embryos (Figure 3H and I). The entropy values are 12.47±0.02 bits in control, while entropy values are 12.55±0.02 bits in Rho1 CA and 12.61±0.02 bits in Rho1 DN.

Next, we assessed the impact of Rho signaling on the mobility of E-cadherin by the FRAP experiments (Figures 3J–M). We observed a doubling of the mobile fractions after either up- or down-regulating Rho signaling in stage 8 embryos (Figure 3L), whereas the half-time remained almost unchanged (Figure 3M). Taken together, our findings are consistent with a model that balanced Rho signaling and, correspondingly, actomyosin contractility boosts E-cadherin junctional recruitment and cluster formation and distribution.

### RhoGEF2 is required for E-cadherin junctional recruitment and immobilization

Rho signaling is controlled by upstream RhoGEFs and RhoGAPs ^43^. Previous studies identified distinct RhoGEFs for activation of medial and junctional actomyosin contractility in gastrulating *Drosophila* embryos, in which RhoGEF2 triggers the medial-apical Rho signaling in the lateral epidermis ^32,40,41^. To analyze whether RhoGEF2 also impacts on E-cadherin, we applied our assays on *RhoGEF2* mutants (embryos from females with *RhoGEF2* mutant germline clones). In fixed and living embryos, we observed reduced medial-apical Myo-II and E-cadherin levels at junctions in *RhoGEF2* mutant embryos (Figures 4A, B and S3) consistent with previous studies ^39,40^. The mean fluorescence intensity of E-cadherin at junctions was half in mutants (Figures 4C and S3B). Meanwhile, the total protein amounts assayed by western blot were comparable to wild-type embryos (Figures 3F and G). We further examined the distribution and mobility of E-cadherin-GFP in *RhoGEF2* mutant embryos. The clusters apparent in high-resolution images were rarely detectable in the high-resolution images from the *RhoGEF2* mutant embryos (Figure 4D). Accordingly, we found an entropy value (12.59±0.02 bits) in the *RhoGEF2* mutant higher than in wild-type embryos (12.42±0.02 bits) (Figure 4E). Consistent with reduced clustering, the mobile fraction was larger in *RhoGEF2* mutants than in wild-type embryos (Figures 4F–I). Thus, our data support a model that RhoGEF2, at least in part, controls Rho signaling, actomyosin contractility, and E-cadherin at junctions in gastrulating embryos.

**Figure 4.**
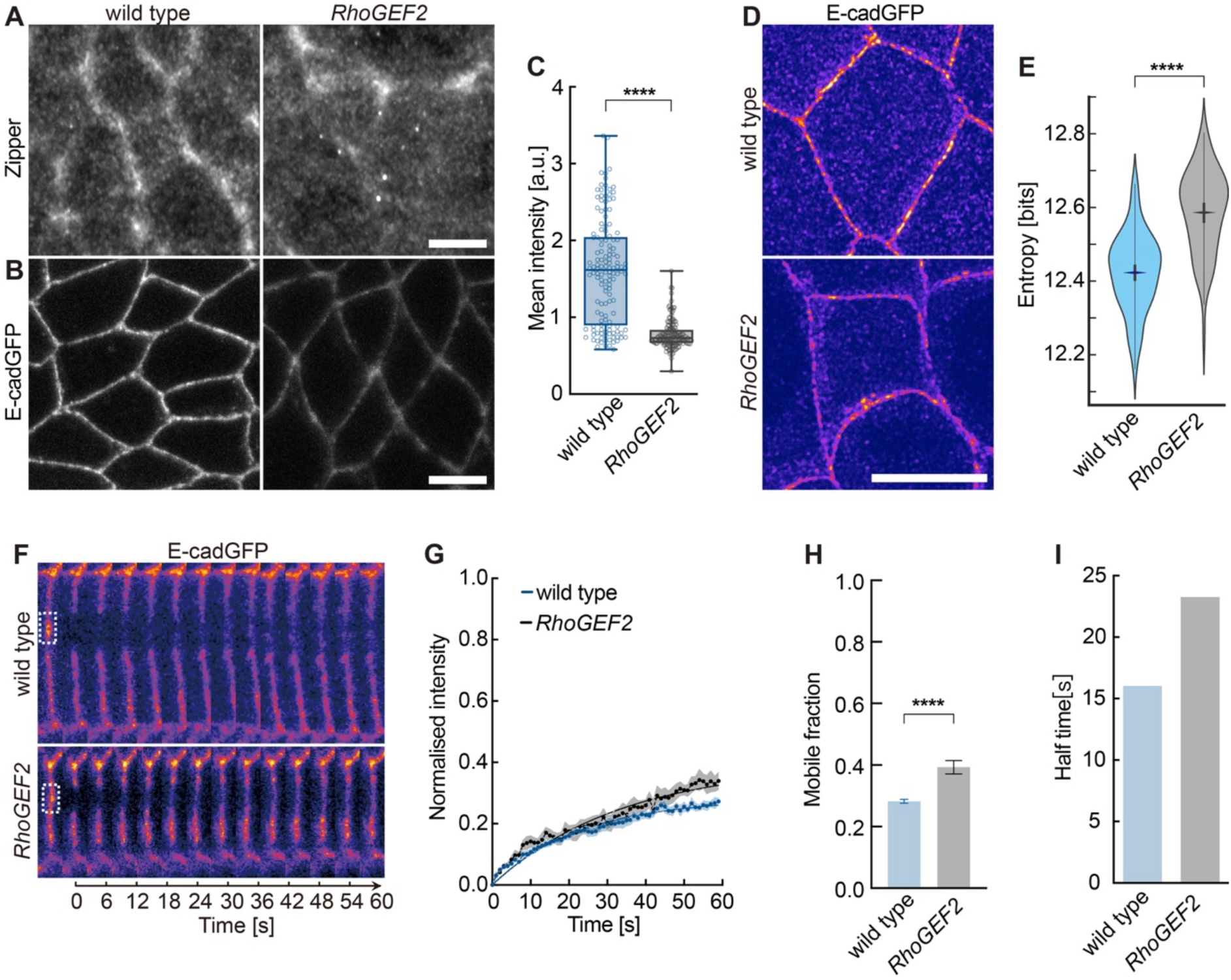
RhoGEF2 is required for E-cadherin recruitment and distribution at cell junctions. **(A, B)** Representative images of Myosin-II (detected by Zipper antibody) **(A)** and E-cadherin-GFP **(B)** from wild-type and *RhoGEF2* germline clone (*RhoGEF2* mutant) embryos. **(C)** Quantifications show junction E-cadherin recruitment reduced in *RhoGEF2* mutant embryos. N = 138 (control) and 162 (*RhoGEF2*) junctions from 5 embryos in each genotype. **(D)** Representative high-resolution images of E-cadherin-GFP by Airyscan jDeconvolution from wild-type and *RhoGEF2* mutant embryos. **(E)** Distribution of entropy values. The rhombus designates the median entropy value; thick bars indicate 95% confidence intervals, and the width is a kernel density estimate of the underlying distribution. Each entropy value was calculated by a 2.2 µm (200 pixels) long junction patch segment with a 0.44 µm (40 pixels) width. **(F-I)** E-cadherin-GFP fluorescence recovery after photobleaching in wild type versus *RhoGEF2* mutant embryos. **(F)** Kymograph of E-cadherin-GFP pre- and post-bleaching, **(G)** normalized fluorescence intensity, **(H)** E-cadherin-GFP mobile fractions, and **(I)** mean recovery times after photobleaching. N = 11 (wild type) and 8 (*RhoGEF2*) junctions from more than five embryos from each genotype. The boxes extend in C from the 25^th^ to the 75^th^ percentiles and the whiskers to the minimum and the maximum value with individual dots. Data are mean ± SEM in G and H. The unpaired two-sided t-test estimated p-values, ns > 0.05, *** p < 0.001, **** p < 0.0001. Scale bars, 5 μm.

### Medial Myosin-II promotes E-cadherin junctional recruitment but not the immobilization

Having analyzed a loss-of-function for *RhoGEF2*, we also aimed for a gain-of-function situation, i. e., increased activation of Rho signaling. As RhoGEF2 overexpression by a maternal GAL4 driver impairs cellularization, we employed the upstream factor Gα12/13. Consistent with a previous report ^39^, medial but not junctional Myo-II levels were increased in embryos overexpressing Gα12/13 (Figures 5A, C, D). Correspondingly, E-cadherin-GFP was increased by more than 50% in Gα12/13 overexpressing embryos compared to control embryos (Figures 5B and E) at cell junctions. Furthermore, the gain-of-function situation also affected the distribution of E-cadherin clusters based on high-resolution images obtained with AiryScan recording and joint deconvolution (Figure 5F). We calculated a higher entropy in Gα12/13 overexpressing (12.53±0.01 bits) than in control embryos (12.47±0.02 bits) (Figure 5G). Consistent with the increased entropy value, we detected a significantly increased mobile fraction in FRAP experiments in the Gα12/13 overexpression embryos (Figure 6H–K). Thus, we presented evidence that the previously reported upstream regulators of Rho signaling and cortical Myo-II impinge on E-cadherin clustering.

**Figure 5.**
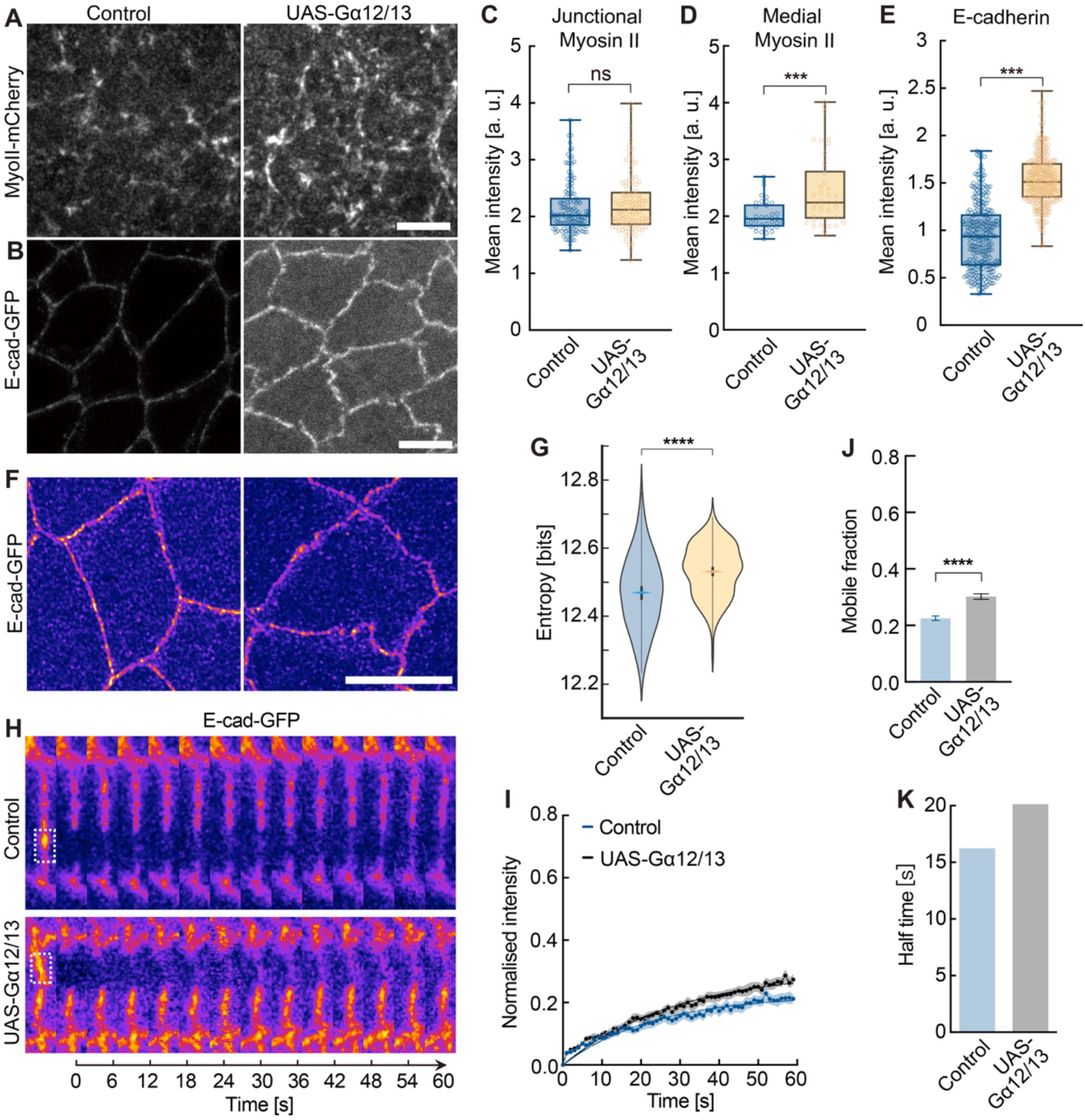
Medial Myosin-II tunes E-cadherin recruitment, distribution, and immobilization at cell junctions. **(A, B)** Representative images of Myosin-II-mCherry **(A)** and E-cadherin-GFP **(B)** from control and Gα12/13 overexpressed embryos. **(C)** Quantifications show no significant difference in the junctional Myosin-II between the control and Gα12/13 overexpressed embryos, N = 133 (control) and 102 (Gα12/13) junctions from 3 embryos in each genotype. **(D)** Quantifications show Medial Myosin-II increased in Gα12/13 overexpressed embryos, N = 37 (control) and 38 (Gα12/13) cells from 3 embryos in each genotype. **(E)** Quantifications show junction E-cadherin recruitment increased in Gα12/13 overexpressed embryos, N = 298 junctions from 5 embryos in each genotype. **(F)** Representative high-resolution images of E-cadherin-GFP by Airyscan jDeconvolution from control and Gα12/13 overexpressed embryos. **(G)** Distribution of entropy values. The rhombus designates the median entropy value; thick bars indicate 95% confidence intervals, and the width is a kernel density estimate of the underlying distribution. Each entropy value was calculated by a 2.2 µm (200 pixels) long junction patch segment with a 0.44 µm (40 pixels) width. Scale bars, 5 μm. **(H-J)** E-cadherin-GFP fluorescence recovery after photobleaching in control versus Gα12/13 overexpressing embryos. **(H)** Kymograph of E-cadherin-GFP pre- and post-bleaching, **(I)** normalized fluorescence intensity, **(J)** E-cadherin-GFP mobile fractions, and **(K)** mean recovery times after photobleaching. N = 16 (control) and 15 (Gα12/13) junctions from more than five embryos in each genotype. The boxes extend in C-E from the 25^th^ to the 75^th^ percentiles and the whiskers to the minimum and the maximum value with individual dots. Data are mean ± SEM in I and J, and solid curves indicate fitting in I. The unpaired two-sided t-test estimated p-values, ns > 0.05, *** p < 0.001, **** p < 0.0001. Unpaired two-sided *t-test* estimates the p-values, **** p < 0.0001.

**Figure 6.**
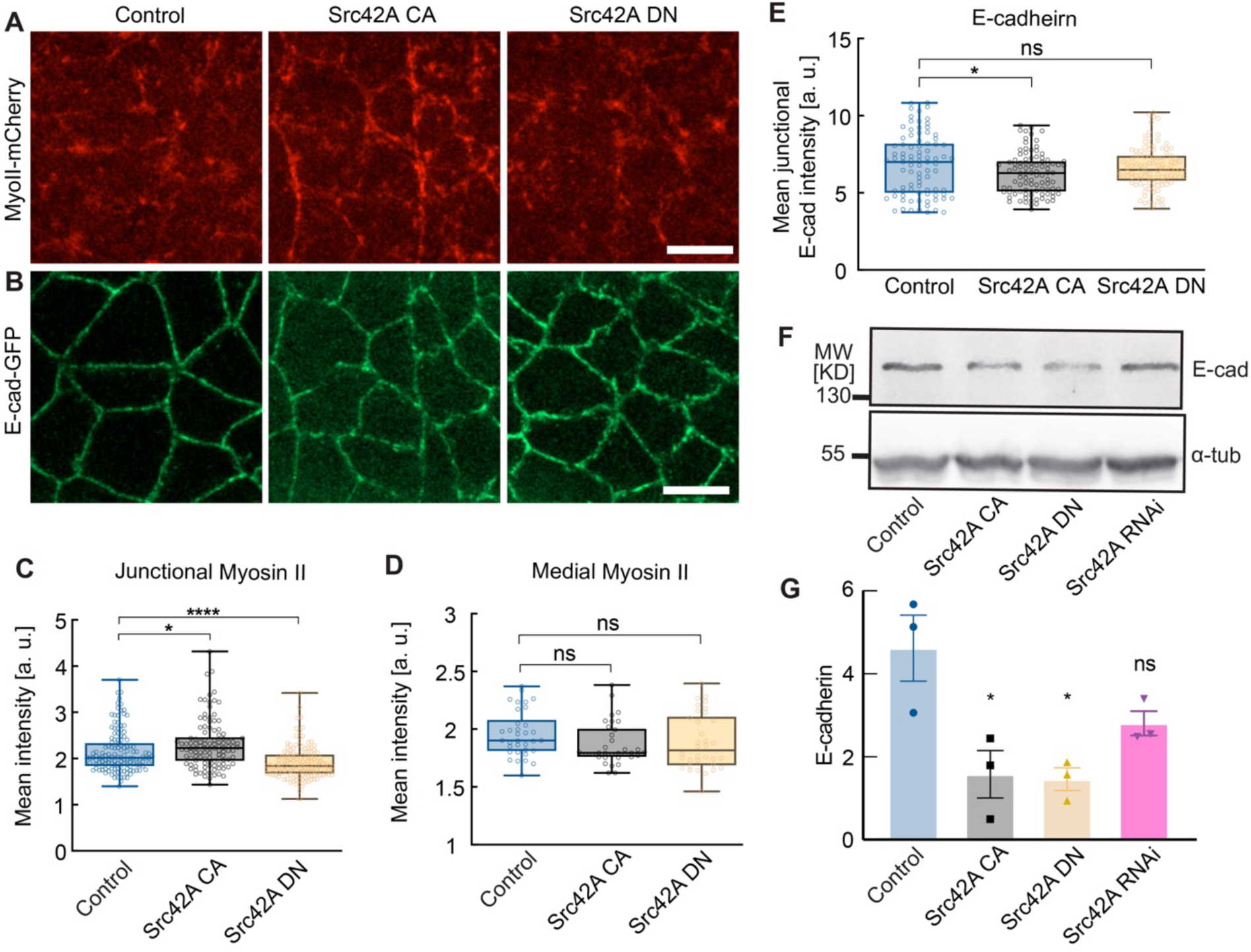
Junctional Myosin-II via Src42A tunes E-cadherin recruitment at cell junctions. **(A, B)** Representative images of Myosin-II-mCherry **(A)** and E-cadherin-GFP **(B)** from control, Src42A CA (constitutively active Src42A), and Src42A DN (dominant negative Src42A) embryos. **(C, D)** Quantifications show the junctional and medial Myosin-II were altered in Src42A CA and Src42A DN embryos. N = 133 (control), 110 (Src42A CA), and 142 (Src42A DN) junctions **(C)**; N = 34 (control), 34 (Src42A CA), and 34 (Src42A DN) cells **(D)** from 3 embryos in each genotype. **(E)** Quantifications show junction E-cadherin recruitment alters in Rho1 CA and Rho1 DN embryos. N = 87 (control), 108 (Src42A CA), and 96 (Src42A DN) junctions from 3 embryos in each genotype. **(F, G)** No significant difference in the E-cadherin total protein was detected from Rho1 mutants compared to control embryos. **(F)** Western blot of total embryonic extracts (3-6 h) with E-cadherin antibody, phenotypes as indicated. Loading was controlled by western blot with α-tubulin antibody. **(G)** Quantification of E-cadherin protein level relative to α-tubulin from the western blot of total embryonic extracts, phenotypes as indicated. The boxes extend (in C-E) from the 25^th^ to the 75^th^ percentiles and the whiskers to the minimum and the maximum value with individual dots. The bar graph indicates the mean, the error bars indicate the SD, and each dot indicates one western blot. An unpaired two-sided t-test estimated p-values, ns > 0.05, * p < 0.05. Unpaired two-sided *t-test* estimates the p-values, ns > 0.05, * p < 0.05, **** p < 0.0001. Scale bars, 5 μm.

### Junctional Myosin-II suppresses E-cadherin junctional recruitment

The protooncogene Src is a major regulator of adherens junctions, as demonstrated in colon cancer cells, for example ^10,44^. In Drosophila embryos, *Src* genetically interacts with E-cadherin and regulates E-cadherin dynamics ^45–48^. In parallel to Rho signaling, Src controls Myo-II levels at junctions in the gastrulating embryos ^25^. We investigated whether Src is involved in the developmental maturation of adherens junctions between stages 6 and 8, besides the previously demonstrated general role. We experimentally interfered with Src function by expressing either a constitutively active version of Src42A (Src42A CA) or a dominant negative version of Src42A (Src42A DN) ^46,48^. Consistent with a previous study ^25^, Myo-II levels at junctions were increased by Src42A CA, while suppressed by Src42A DN as assayed in fixed in live embryos (Figures 6A, C and S4A–F). In contrast, we did not detect changes in medial Myo-II (Figure 7D). Next, we examined the E-cadherin levels in these embryos and found a weak but significant reduction in E-cadherin-GFP levels in Src42A CA embryos but not in Src42A DN embryos (Figures 7B, E and S4G). Interestingly, we measured decreased total E-cadherin levels in Src42A CA and Src42A DN embryos by western blots, which may be caused by increased endocytosis and turnover of E-cadherin.

**Figure 7.**
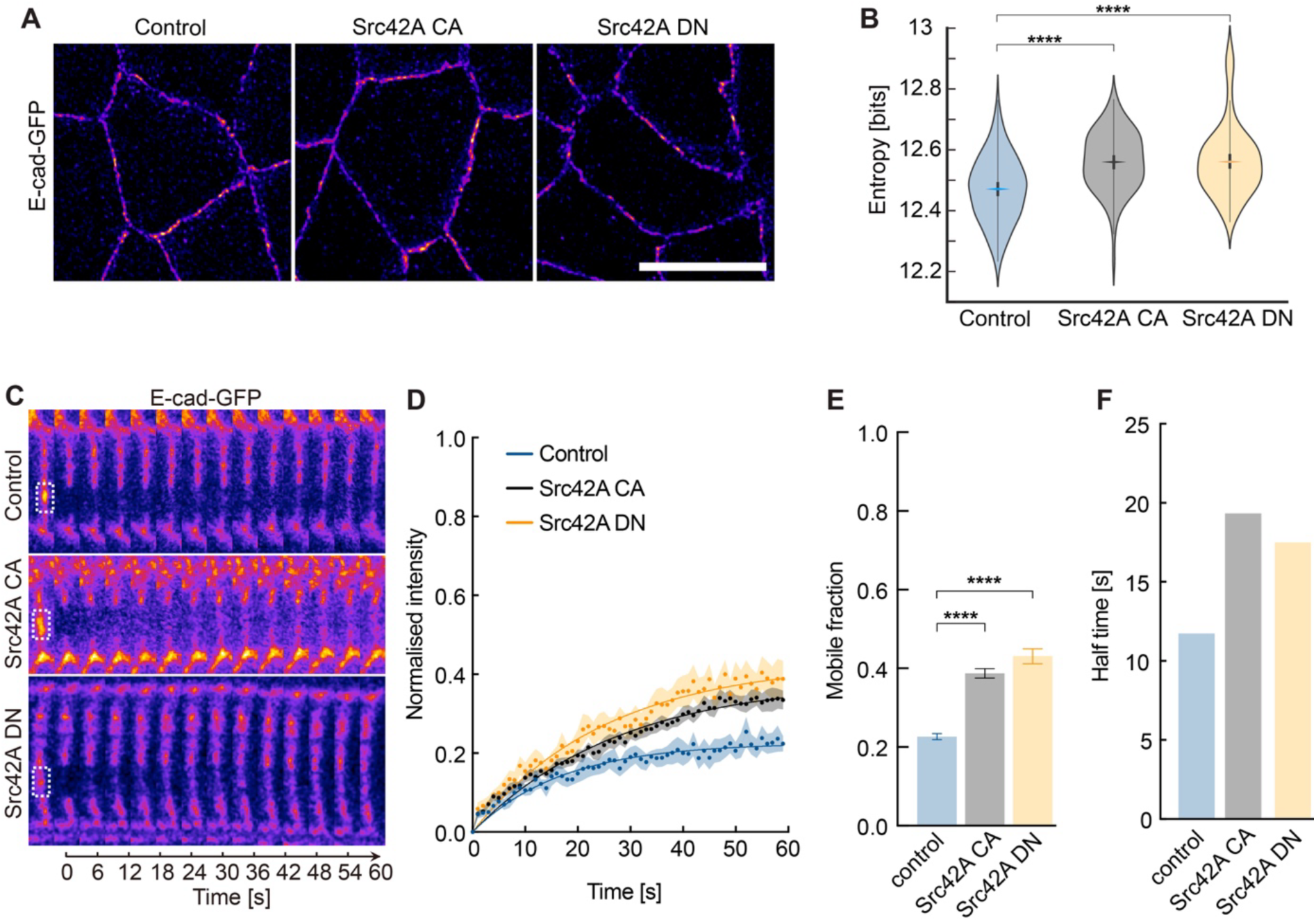
Src42A alters the distribution and immobilization of E-cadherin. **(A)** Representative high-resolution images of E-cadherin-GFP by Airyscan jDeconvolution from control, Src42A CA, and Src42A DN embryos. **(B)** Distribution of entropy values. The rhombus designates the median entropy value; thick bars indicate 95% confidence intervals, and the width is a kernel density estimate of the underlying distribution. Each entropy value was calculated by a 2.2 µm (200 pixels) long junction patch segment with a 0.44 µm (40 pixels) width. **(C-F)** E-cadherin-GFP fluorescence recovery after photobleaching in control versus Src42A CA and Src42A DN embryos. **(C)** Kymograph of E-cadherin-GFP pre- and post-bleaching, **(D)** normalized fluorescence intensity, **(E)** E-cadherin-GFP mobile fractions, and **(F)** mean recovery times after photobleaching. N = 8 (control), 12 (Src CA), and 10 (Src DN) from more than five embryos in each genotype. Data are mean ± SEM in (D and E), and solid curves indicate fitting in (D). Unpaired two-sided *t-test* estimates the p-values, **** p < 0.0001.

Consistent with the impact of Src signaling on E-cadherin levels, we detected changes in E-cadherin distribution and mobility following up- and down-regulation of Src. Quantification of high-resolution images revealed an increased entropy by both Src42A CA (12.56±0.02 bits) and Src42A DN (12.56±0.02 bits) compared to the control (12.47±0.02) (Figure 7A and B). Similar to the changed distribution pattern of clusters, the mobile fractions were increased and thus closer to the situation in stage 6 embryos: 38.7%±1.2 with Src42A CA and 43.1%±1.9 with Src42A DN (Figure 7C–F). We also detected longer recovery time in Src42A CA/DN than in control embryos (Figure 7F). Depletion of Src42A by RNAi led to comparable data (Figure S5). Thus, our data suggests that Src42A impinges on the developmental control of E-cadherin during gastrulation, probably by modulating the junctional Myo-II levels.

## DISCUSSION

A fundamental characteristic of metazoan development and epithelial tissue is stable intercellular cohesion. The distribution and molecular dynamics of the adhesion receptors are governed precisely from cell-cell contact formation to the establishment of the matured junction ^31^. Our study established an assay for the developmentally controlled maturation of AJs in the epithelial cells of *Drosophila* embryonic ectoderm during gastrulation. Our data revealed the type and degree of E-cadherin changes in total amount, junctional recruitment, and clusters’ immobilization. We combined high-resolution imaging and entropy calculation to assay the pattern of E-cadherin distribution along the junctions, which revealed that the certainty about the location of E-cadherin clusters.

E-cadherin levels and turnover at AJs have previously been shown to be controlled by actomyosin contractility ^6,7,39,49,50^. The junctional accumulation and reduced turnover of E-cadherin occur parallel to Myo-II levels increasing during gastrulation. We show a requirement of contractility by ROCK inhibitor injection for E-cadherin junctional levels in stage 8, consistent with a previous study ^39^. Also consistent with results from a study with cultured cells ^6^, the mobile fraction of the remaining E-cadherin at the cell junctions increased threefold, comparable to the stage 6 embryos. Thus, consistent with the AJs maturation in the mammalian cell, actomyosin contractility controlled the developmental maturation of AJs in gastrulating embryos.

It has been reported previously that actomyosin contractility is regulated by Rho signaling and Src kinase in gastrulating embryos ^23–25^. However, how these components control the developmental maturation of AJs has remained unclear. Our data suggest that Rho signaling and Src kinase have distinct and shared roles in AJs’ maturation. Overall Rho up- or down-regulating increases or decreases E-cadherin levels at junctions, respectively. We also identify upstream regulators of Rho signaling controling the E-cadherin junctional levels. E-cadherin levels at junctions were reduced in *RhoGEF2* mutant embryos while increased in the Gα12/13 overexpression embryos. In several of the above situations, the total protein amount is not affected as expected. Thus, our data suggest that Rho signaling promotes E-cadherin junctional recruitment during gastrulation.

Rho signaling also controls the turnover of E-cadherin clusters ^51–53^. As expected, the mobile fraction of E-cadherin-GFP increased after down-regulation of Rho signaling or in *RhoGEF2* mutant embryos. Although the junctional level of E-cadherin is increased following Rho upregulation or Gα12/13 overexpression embryos, to our surprise, we also detected a higher mobile fraction in these embryos, suggesting the E-cadherin clustering is regulated in a precision manner. Interestingly, the entropy values are also higher in the embryos detected with a higher mobile fraction, suggesting that the E-cadherin clustering and clusters’ distribution are highly sensitive to mechanical tension. The mobile fraction is lower in the E-cadherin clusters than the relatively uniform distributed E-cadherin both in mammalian cells and *Drosophila* embryos ^6,16^. Our data show both the entropy values and the mobile fraction are lower in the mature junctions. Given the key function of E-cadherin for mechanotransduction ^54,55^, we expect a link between the E-cadherin distribution pattern and immobilization, which remains to be determined by super-resolution imaging, for example.

Our research confirms that Src kinase is a key regulator of Myo-II in gastrulating embryos besides the Rho signaling ^25^. The expression of either Src42A CA or DN in the embryos up- or down-regulates the junctional rather than the apical-medial Myo-II, providing robust evidence for this regulatory role. Although the Src42A DN embryos do not affect the E-cadherin junctional level, we revealed that junctional recruitment is suppressed in the Src42A CA embryos. Our data provide experimental evidence for a model-predicted idea that the shear forces decrease the E-cadherin level ^39^. Src kinase also controls cellular adhesions in the mammalian cell ^56^. It co-localizes with E-cadherin at the cell-cell adhesion in non-migrating epithelial cells ^57^; however, Src activity leads to the release of β-catenin from AJs and suppresses E-cadherin in cancer cell epithelial-to-mesenchymal transition ^58,59^, targets lysosomal degradation of E-cadherin ^60^, and induces the endocytosis of E-cadherin to disrupt cell-cell contacts in the MDCK cells ^61^. We also detected a slight decrease in the total E-cadherin protein level in Src42A CA and DN embryos. It remains to be investigated in future experiments whether Src also controls lysosomal degradation of E-cadherin in embryos. Nevertheless, our data suggest that Src kinase regulates E-cadherin developmental maturation in gastrulating embryos.

Similar to our experiments about the contribution of Rho signaling, when perturbing Myo-II activity by Src42A CA or DN, E-cadherin distribution and turnover were likewise influenced. Either Src42A CA or DN expression increases the entropy values and the mobile fraction, which further supports the idea that E-cadherin clustering and clusters’ distribution are highly sensitive to mechanical tension.

In summary, our findings characterize the developmental control of E-cadherin in cells of the lateral epidermis during gastrulation (Figure S6). Both Rho signaling and Src mediate E-cadherin at cell junctions and regulate its distribution and immobilization. More generally, these findings provide that the mechanical microenvironment under E-cadherin molecules controls its molecular dynamics and clustering to achieve precision regulation in a self-organized manner during tissue morphogenesis.

## MATERIALS AND METHODS

### Drosophila genetics

The following transgenes or mutants are used in this study. If not otherwise noted, stocks were generated in this study or obtained from the Bloomington Drosophila Stock Centre ^62^ and others. A FlyBase Symbol ^63^ is provided here. OrR; His-GFP (w; Histone2Av-GFP) (Bloomington number 24163) is GFP-Histon2Av expressed with a genomic construct; *RhoGEF2* (w; Frt[G13]{w+} l(2)04291{ry+}/CyO) is a P allele of RhoGEF2, apparently no transcript of RhoGEF2 ^64^; *UAS-Rho^V^*^14^ (Rho1 CA, constitutively active Rho1) (Bloomington number 7330, Flybase Symbol, Dmel\P{UAS-Rho1.V14}5.1, a gift from Natalia A. Bulgakova) ^53^; *UAS-Rho^N^*^19^ (Rho1 DN, dominant negative Rho1) (Bloomington number 7328, Flybase Symbol, Dmel\P{w[+mC]=UAS-Rho1.N19}2.1, a gift from Natalia A. Bulgakova) ^53^; *UAS-Src42A^CA^* (Src42A CA, constitutively active Src42A, w; Sp/CyO Dfd-GMR-nvYFP; P{w[+mC]=UAS-Src42A.CA}5, a gift from Stefan Luschnig) ^46,65^; *UAS-Src42A^DN^*(Src42A DN, dominant negative Src42A, w; UAS-Src42A.DN (K295M)/TM6b Dfd-GMR-nvYFP, a gift from Stefan Luschnig) ^46,48^; *UAS-Gα12/13* (a gift from Thomas Lecuit) produces wild-type Gα12/13, the α-subunit of the heterotrimeric G-protein complex that associates with GPCR *smog* ^66^. *matGal67-15* (w; tub-Gal4-VP16{w+}[67]; tub-Gal4-VP16{w+}[15]) is a ubiquitous, maternally supplied, Gal4 driver (a gift from Daniel St Johnston).

E-cadherin-GFP (E-cadherin-GFP) (named as wild-type in this study) is a homozygous viable DE-cadherin knock-in at the locus ^28^. It is used to quantify the E-cadherin recruitment at cell junctions in Figures 1B–D, 2A, 2B, 3B, 3E, 4B, 4C, 5B, 5E, 6B, and 6E; exemplify E-cadherin distribution in Figures 1K, 1L, 3H, 3I, 4D, 4E, 5F, 5G, 7A and 7B; investigate the E-cadherin turnover by FRAP in Figures 1G–J, 2C–F, 3J–M, 4F-I, 5H–K, 7C–F, and S5.

To investigate the E-cadherin recruitment, distribution, and turnover at cell junctions under different tension-altering conditions, “+; 67-Gal4, Myo-II-mCherry, E-cadherin-GFP;+” (a gift from Thomas Lecuit) ^39^ was applied, in which the E-cadherin-GFP, 67-Gal4, and Myosin-II-mCherry are recombined. Here, *Myosin-II-mCherry* (Myo-II-mCherry) is a tagged construct of *Drosophila* “non-muscle Myosin-II regulatory light chain” encoded by gene *spaghetti squash* downstream of its native ubiquitously active promoter ^67^; *67-Gal4* (mat αTub-GAL4-VP16) is a ubiquitous, maternally supplied, Gal4 driver.

To alter the tension conditions in the embryos, “+; 67-Gal4, Myo-II-mCherry, E-cadherin-GFP;+” virgin females were collected and crossed with males with different genotypes separately: *y, w* (control), *UAS-Rho^V^*^14^ (Rho1 CA), *UAS-Rho^N^*^19^ (Rho1 DN), *UAS-Gα12/13* (Gα12/13 over-expression), *UAS-Src42A^CA^*(Src42A CA), *UAS-Src42A^DN^*(Src42A DN). F1 virgin females were collected and crossed with E-cadherin-GFP males for embryo collection.

To detect the endogenous E-cadherin expression in the embryos (Figures 3F, 3G, 6E, and 6F), *matGal67-15* virgin females were collected and crossed with males with different genotypes separately. F1 virgin females were collected and crossed with OrR males for embryo collection.

*RhoGEF2* was used alone (Figure 3F, 3G, and S3) or recombined with E-cadherin-GFP (Figure 4) to generate the germ-line clone. *RhoGEF2* germline clones were generated and selected with *ovo^D^*. The first and second instar larvae were heat-shocked twice for 60 min at 37℃, and virgin females were collected and crossed with OrR or E-cadherin-GFP males for embryo collection.

E-cadherin turnover analysis in Src42A RNAi, y1 w*; P{matα4-GAL4-VP16}67 E-cadherin-GFP; P{matα4-GAL4-VP16}15/ P{TRiP.HMC04138}attP2 flies were crossed to obtain Src42A RNAi embryos along with an E-cadherin-GFP marker ^47^.

### Immunostaining and imaging

2.5-5 hours embryos were fixed with 4% formaldehyde. Endogenous E-cadherin was stained by DCAD2 antibody (rat, 7 μg/ml) ^68^ in Figures S3. His-GFP embryos were stained in the same tube as the control. A GFP nanobody (GFP-Booster ATTO488, 1:250, Chromo Tek) was used to detect the E-cadherin-GFP in high-resolution images (Figures 1K, 3H, 4D, 5F, and 7A). For the Myosin-II staining, 2.5–5 hours embryos were heat-fixed in a salt solution (0.4% NaCl, 0.03% Triton X-100). The Myosin-II heavy chain was detected by an anti-Zipper polyclonal antibody (Rabbit anti-Zipper, 1:1000) ^34^, the cell outline was visualized by anti-Neurotactin (BP106, 1:20) staining. Secondary antibodies were labeled with Alexa dyes (Thermo Fisher Scientific, 5 μg/ml). Embryos were mounted in Aquapolymount (Polysciences) for imaging. The apical plane of the embryo was acquired with axial sections of each 0.5 µm on a laser scanning confocal microscopy (Carl Zeiss, ZEISS LSM 980 with Airyscan 2) with a 63x oil objective (Carl Zeiss, 63x/oil, NA1.4).

### High-resolution imaging and entropy analysis

High-resolution images (Figures 1K, 3H, 4D, 5F, and 7A) were acquired with axial sections of each 0.17 µm by Airyscan 2 detection (ZEISS LSM 980 with Airyscan 2, Carl Zeiss) with a 63x oil objective (63x/oil, NA1.4, Carl Zeiss) ^69^. The resolution was improved by joint Deconvolution (jDeconvolution) processing ^70^. The pixel size is 0.011 μm. 8-10 image stacks were processed with “max intensity” in ImageJ/ Fiji, and the processed images were segmented using the “Tissue Analyzer” ^71^, a plugin in ImageJ/ Fiji ^72^. The vertices (tricellular junctions) were avoided, and the individual cell junctions were selected with 41-pixel widths to straighten by the “straighten” tool in ImageJ/ Fiji for the entropy analysis.

The (discrete) Shannon entropy, the *S* = ∑ *p*_*i*_log_2_*p*_*i*_, where *p_i_ is* normalized frequencies, serves as a measure of uncertainty in information theory and statistical physics ^29,30^. It depends on the shape of a probability distribution where unimodal sharp distributions yield lower values and uniform distributions or distributions with many peaks yield high entropy values. The Shannon entropy of each 2.2 μm x 0.44 μm patch along junctions by treating the E-cadherin intensity signal as a two-dimensional probability distribution. Intensities were normalized to obtain a “probability” of the E-cadherin locations *p*_*i*_ = *I*_*i*_/ ∑_*i*_ *I*_*i*_ where the index *i* runs over all pixels in the patch.

### Live imaging

Embryos were prepared as previously described ^73^. Briefly, staged embryos were collected and dechorionated with 50% hypochlorite bleach for 90 seconds, aligned on an agar block, and attached to the coverslips by homemade glue covered with halocarbon oil.

For the quantification of E-cadherin recruitment at cell junctions, images were acquired on a laser scanning confocal microscopy (ZEISS LSM 980 with Airyscan 2, Carl Zeiss) with a 63x oil objective (63x/oil, NA1.4, Carl Zeiss) under the confocal model with a unit pinhole (0.51 μm). The apical plane of the embryo was acquired with axial sections of each 0.5 µm. The image size is 33.67 x 33.67 μm (512 x 512 pixels).

For the quantification of Myosin-II, image stacks with axial sections of each 0.5 µm were acquired by Airyscan 2 detection on a laser scanning confocal microscopy (ZEISS LSM 980 with Airyscan 2, Carl Zeiss) with a 63x oil objective (63x/oil, NA1.4, Carl Zeiss). E-cadherin-GFP channel was used as a reference for cell junctions and acquired in parallel. The image size is 26.43 x 26.42 μm (620 x 620 pixels).

### Fluorescence recovery after photobleaching (FRAP)

Embryos were prepared as previously described ^38^. Briefly, staged embryos were collected and dechorionated with 50% hypochlorite bleach for 90 seconds, aligned on an agar block, and attached to the coverslips by homemade glue covered with halocarbon oil. A cross-section was acquired using laser scanning confocal microscopy (ZEISS LSM 980 with Airyscan 2, Carl Zeiss) with a 63x oil objective (63x/oil, NA1.4, Carl Zeiss) and a frame rate of 1 Hz for 120 seconds. To cover the junctional E-cadherin over recording, the pinhole was set to 1 μm. The image size is 33.67 x 33.67 μm (512 x 512 pixel). The bleaching region of interest (ROIs) for bleaching was for E-cadherin clusters with 0.5 x 1 μm. The ROIs were bleached by 50% of 488 laser intensity with 20 iterations at a scan speed of 5 after the first 2 frames were recorded.

The ROI was tracked to quantify E-cadherin-GFP recovery after photobleaching, and the fluorescence intensity of ROIs was measured manually in Fiji/Image J. The fluorescence intensity was normalized in each recording by

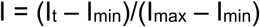

Where,

I_t_ represents the intensity at time t; I_min_ represents the intensity of the first frame after bleaching; I_max_ represents the intensity of one frame before bleaching.

The normalized fluorescence intensity after bleaching was fit by “One-phase association” in GraphPad Prism 10 ^74^, and the mobile fraction and half-time were calculated.

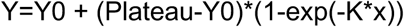

Where,

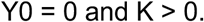

### Y-27632 treatment

Embryos were prepared as previously described ^38^. Embryos were collected and dechorionated with 50% hypochlorite bleach for 90 seconds, dried in a desiccation chamber for ∼10 min, and covered with halocarbon oil. Water (as control) or 20 mM Y-27632 (Sigma) in water was injected dorsally into the vitelline space. After injection, Living imaging and FRAP of E-cadherin-GFP were performed as stated above.

### Image analysis

The image stacks were processed with “max intensity” in ImageJ/Fiji. To quantify E-cadherin recruitment at cell junctions, the processed images were segmented using the “Tissue Analyzer.” The segmentation was used to restrict the ROI of cell junctions; meanwhile, the vertices (tricellular junctions) were excluded. The mean intensity from each junction is measured. To quantify the junctional Myosin-II levels from the anti-Zipper antibody staining in Figueres 4A, S1, and S4, the segmentation was performed from the channel of Neurotactin staining using the “Tissue Analyzer” The ROI of cell junction was restricted, where the vertices were excluded, and loaded to the Myosin-II channel to measure the mean intensity from each junction. To quantify the junctional and medial Myosin-II levels from the live images of Myosin-II-mCherry in Figures 3, 5, and 6, the E-cadherin-GFP channel recorded with Myosin-II-mCherry in parallel was applied for segmentation using the “Tissue Analyzer.” The ROI of junctional and medial-apical masks (Figure S2), where the vertices were excluded, was restricted and loaded to the Myosin-II-mCherry channel. The mean intensity was measured from each junction and cell.

### Immunoblotting

Embryonic extracts were analysed by SDS polyacrylamide electrophoresis and immunoblotting as previously described ^75^. For the analysis of the total protein level of E-cadherin in different stages (Figures 1E and 1F), 2–4 hours E-cadherin-GFP embryos were heat-fixed first in a salt solution (0.4% NaCl, 0.03% Triton X-100). Stage 5, 6, and 8 embryos were collected separately under a stereomicroscope based on morphological features from the pre-fixed embryos and homogenized with a pistil in a 1.5 ml reaction vial in 1× Laemmli buffer and boiled for 10 min. For the western blot in Figures 3F and 6F, 3-6 hours embryos were homogenized with a pistil in a 1.5 ml reaction vial in 1× Laemmli buffer and boiled for 10 min. Proteins were blotted by wet transfer to nitrocellulose filters (100 mA per mini gel, overnight). The blots were incubated with primary antibodies, developed with fluorescently labelled secondary antibodies (LiCOR, 1:20000, 0.05 µg/ml), and recorded with a LICOR system at 16-bit colour depth. E-cadherin was detected by DCAD1 (rat, 1:100, a gift from Tadashi Uemura) ^68^, and tubulin was detected by an anti-α-tubulin antibody (mouse B512, 1:50000, Sigma). For quantification, integrated signals of E-cadherin were normalized by corresponding α-tubulin signals.

### Statistics

Statistical analysis was performed with Prism10. An unpaired two-sided *t*-test estimates the *p* values.

### Compliance and ethics

The author(s) declare that they have no conflict of interest.

## Acknowledgments

We are grateful to Natalia A. Bulgakova, Daniel St Johnston, Yang Hong, Thomas Lecuit, Stefan Luschnig, and Tadashi Uemura for the materials. This work would not be possible without reagents and resources obtained from or maintained by the Bloomington *Drosophila* Stock Centre (NIH P40OD018537), Flybase (MRC grant MR/N030117/1), and Hybridoma Center. This work was in part supported by the Young Taishan Scholars Program of Shandong Province (qnts20191090155), the Natural Science Foundation of China (32070786), and the Deutsche Forschungsgemeinschaft (DFG FOR1756, GR1945/8-2, GR1945/10-1/2, equipment grant INST 160/718-1 FUGG).

## SUPPLEMENTARY DATA

**Figure S1.**
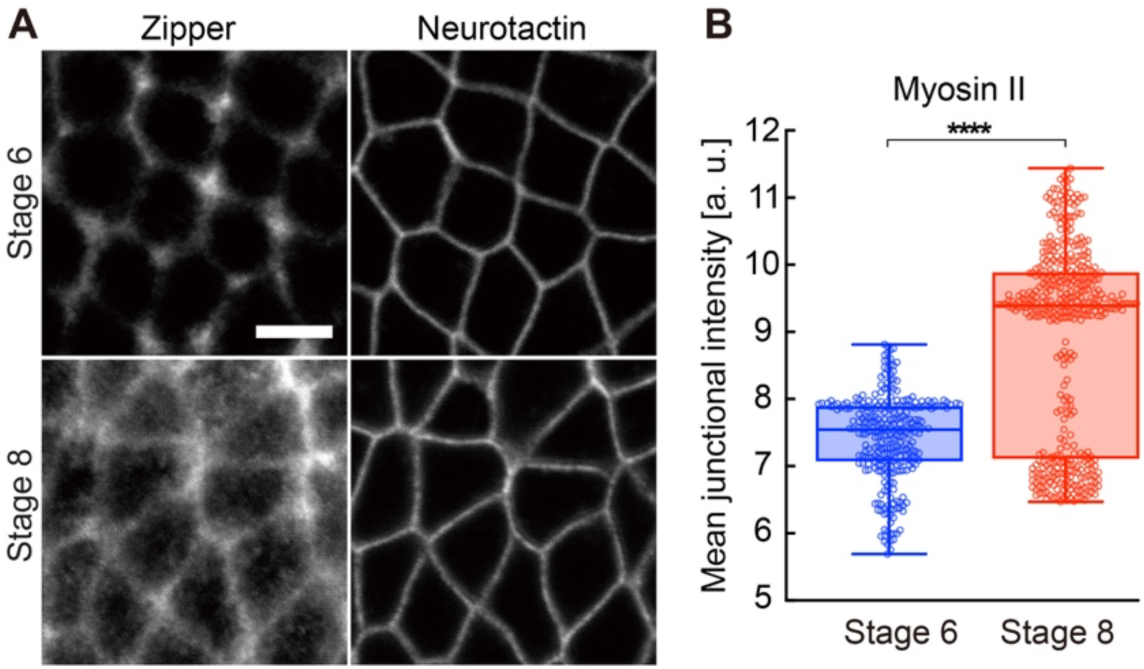
Developmental control of Myosin II. **(A)** Representative images of Myosin-II from the indicated stage embryos. Scale bars 5 µm. **(B)** Quantifications show increased Myosin-II recruitment at cell junction from stage 6 to stage 8. N = 256 junctions from 5 embryos in each stage. The boxes extend from the 25th to the 75th percentiles and the whiskers to the minimum and the maximum value with individual points. Unpaired two-sided t-test estimates the p-values, **** p < 0.0001. Scale bars, 5 µm.

**Figure S2.**
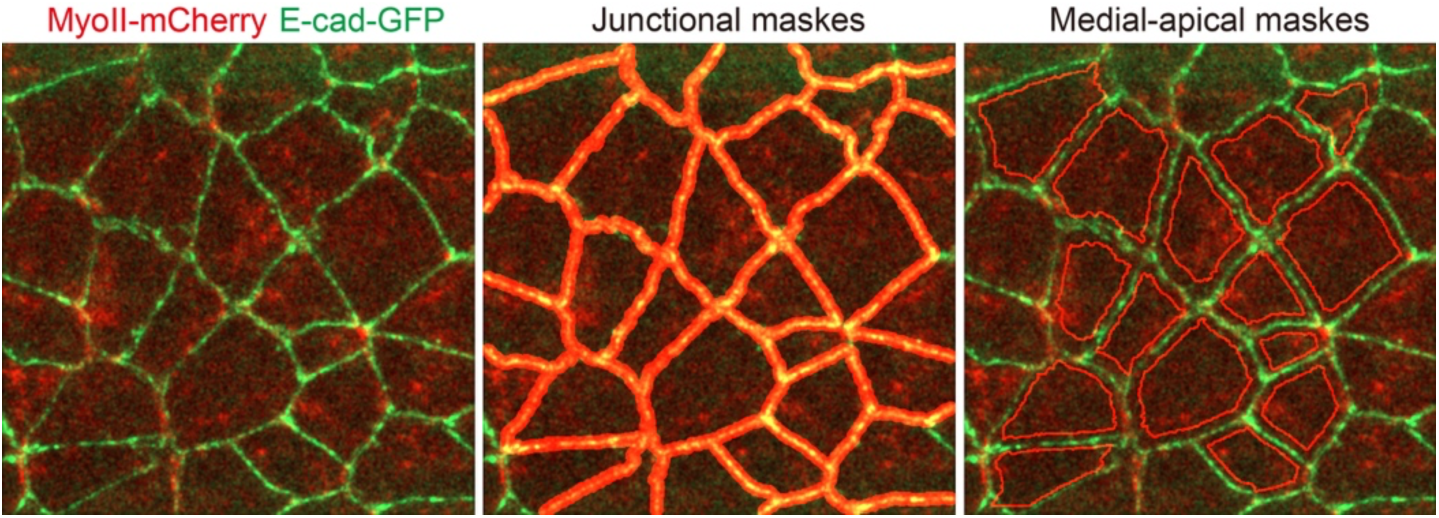
Schematic illustration of Myosin II quantification. Representative images show the junctional and medial-apical masks for the junctional and medical-apical Myosin II intensity measurements.

**Figure S3.**
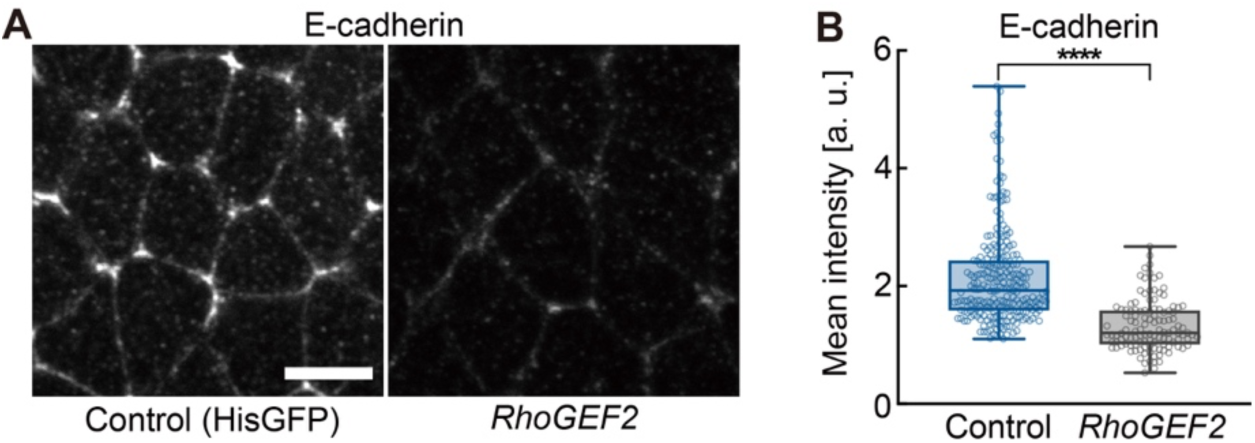
Endogenous E-cadherin junctional recruitment requires RhoGEF2. **(A)** Representative images show the E-cadherin levels from antibody staining in control (Histone-GFP) *RhoGEF2* mutant embryos. **(B)** Quantification of E-cadherin levels at cell junctions. N = 243 (control) and 119 (*RhoGEF2*) junctions from 5 embryos in each genotype. The box extends from the 25th to the 75th percentiles and the whiskers to the minimum and the maximum value with individual points. Unpaired two-sided *t-test* estimates the p-value, **** p < 0.0001. Scale bars 5 µm.

**Figure S4.**
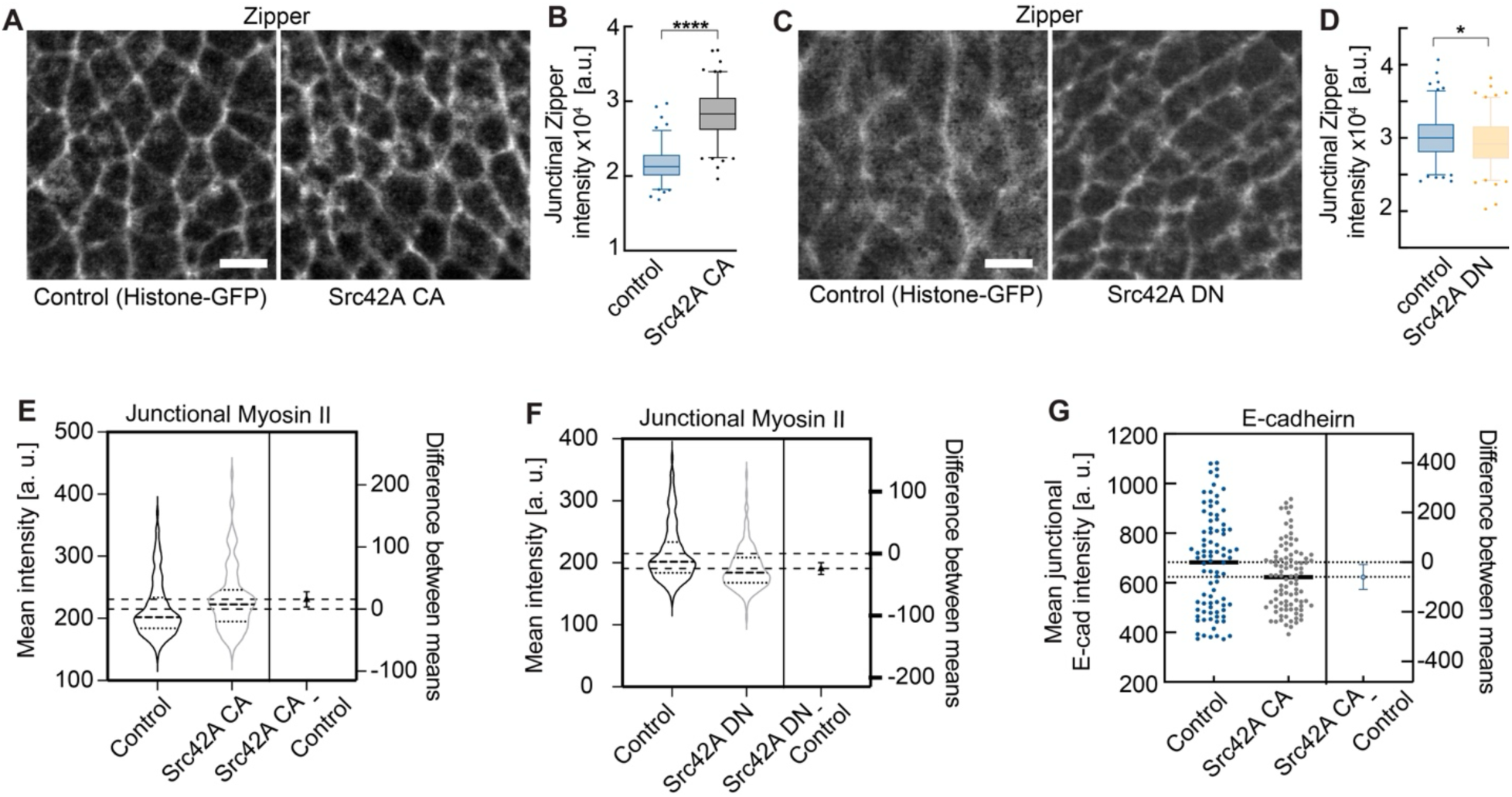
Src42A is required for the endogenous E-cadherin recruitment at cell junctions. **(A, C)** Representative images showing the distribution of Myosin II (anti-Zipper antibody staining) and E-cadherin (anti-E-cadherin antibody staining) in control (Histone-GFP) embryos and embryos expressing Src42A CA or Src42A DN. **(B, D)** Quantification of Myosin II levels at cell junctions. N = 304 junctions from 5 control embryos and 398 junctions from 4 Src42A CA embryos; N = 289 junctions from 3 control embryos and 333 junctions from 3 Src42A DN embryos. The boxes extend from the 25^th^ to 75^th^ percentiles; the whiskers are drawn down to the 2.5^th^ percentile and up to the 97.5^th^; points below and above the whiskers are drawn as individual dots. Unpaired two-sided *t-test* estimates the p-values, * p < 0.05, **** p < 0.0001. **(E, F)** The Difference in quantifying Myosin II levels at cell junctions from the Myo-II mCherry living images between control and Src42A CA (E) or control and Src42A DN (F). **(G) The d**ifference in the quantification of E-cad levels at cell junctions from the E-cad-GFP living images between control and Src42A CA.

**Figure S5.**
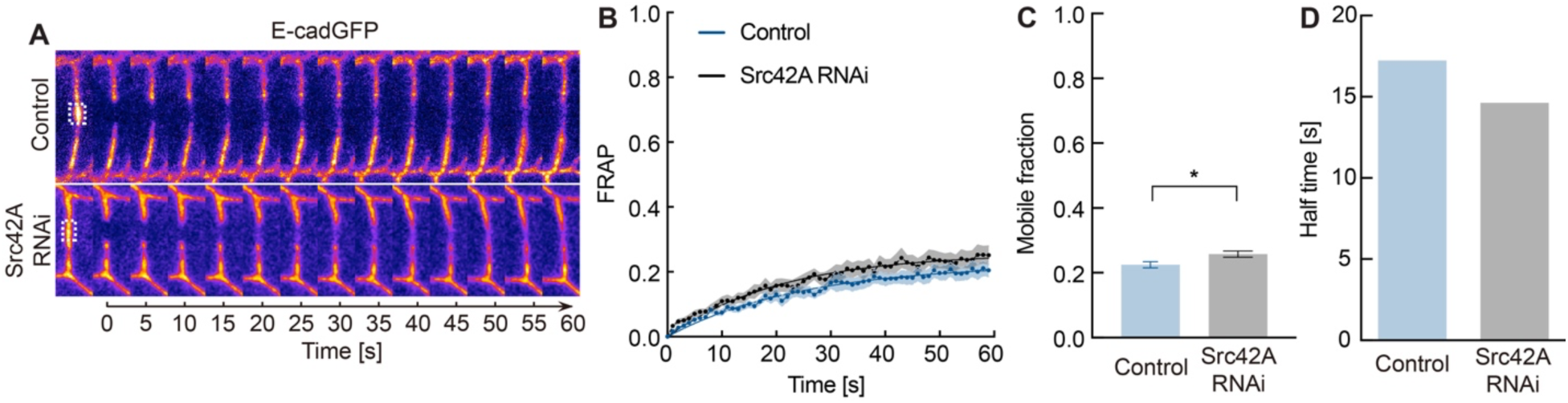
Src42A is required for the E-cadherin immobilization. **(A-D)** E-cadherin-GFP fluorescence recovery after photobleaching in wildtype versus Src42A RNAi embryos. **(A)** Kymograph of E-cadherin-GFP pre- and post-bleaching, **(B)** normalized fluorescence intensity, **(C)** E-cadherin-GFP mobile fractions, and **(D)** mean recovery times after photobleaching. N = 10 from more than five embryos in each genotype. Data are mean ± SEM in (B and C), and solid curves indicate fitting in (B). Unpaired two-sided *t-test* estimates the p-values, * p < 0.05.

**Figure S6.**
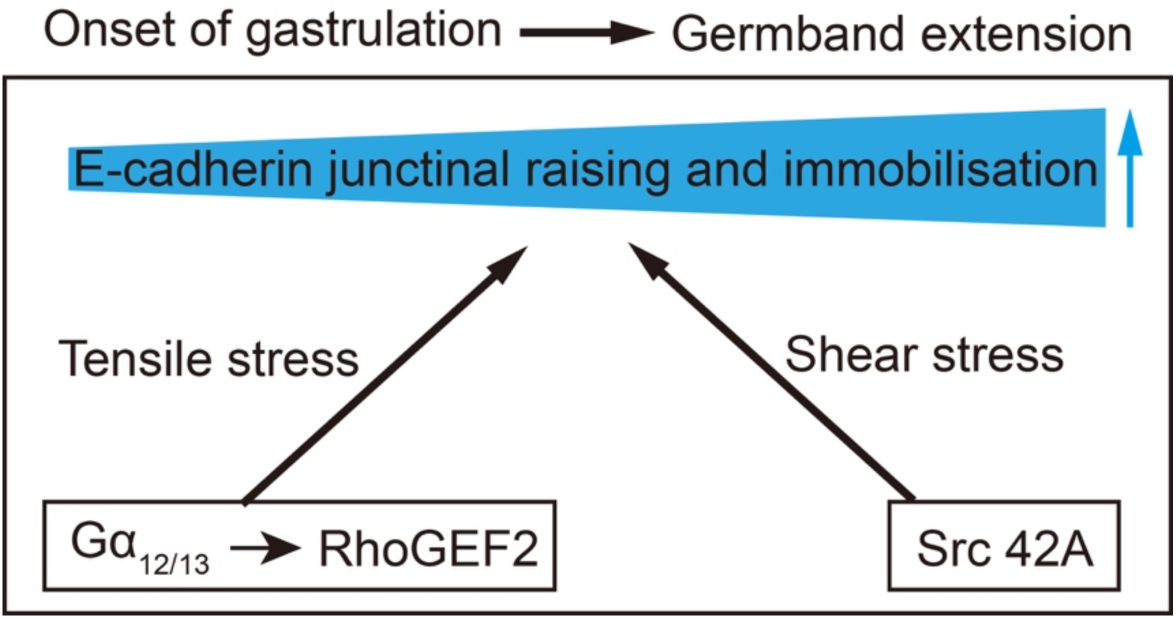
Hypothetical model of the developmental control of E-cadherin at cell junctions.

